# Multiplexed mosaic tumor models reveal natural phenotypic variations in drug response within and between populations

**DOI:** 10.1101/2024.12.13.628239

**Authors:** Johnny X. Yu, Jung Min Suh, Katerina D. Popova, Kristle Garcia, Tanvi Joshi, Bruce Culbertson, Jessica B. Spinelli, Vishvak Subramanyam, Kevin Lou, Kevan M. Shokat, Jonathan Weissman, Hani Goodarzi

## Abstract

Many agents that show promise in preclinical cancer models lack efficacy in patients due to patient heterogeneity that is not captured in traditional assays. To address this problem, we have developed GENEVA, a platform that measures the molecular and phenotypic consequences of drug perturbations within diverse populations of cancer cells at single-cell resolution, both *in vitro* and *in vivo*. Here, we apply GENEVA to study the KRAS G12C inhibitors, recapitulating known properties of these drugs and uncovering a previously unknown role for mitochondrial activation in cell death induced by KRAS inhibition. We demonstrate that this finding can be leveraged for the development of combination therapies with greater efficacy. Finally, we show that the application of GENEVA with *in vivo* mouse models revealed epithelial to mesenchymal transition (EMT) as a key mechanism for resistance to KRAS G12C inhibition.

## Introduction

Targeted therapies aim to precisely attack cancer cells by focusing on their molecular and genetic drivers (*1*). However despite recent advances, variations between and within tumors, known as inter- and intra-tumor heterogeneity respectively, lead to inconsistent responses to these therapies as subpopulations of cancer cells may be inherently more or less sensitive to drugs (*2*). Single-cell genomic technologies, which allow for high-resolution profiling of clinical samples and rich phenotyping of chemical perturbation, have deepened our understanding of cancer cell heterogeneity and drug responses (*3, 4*). However, since these technologies are hard to scale, especially in *in vivo* preclinical models, they are currently absent from the earlier steps in state-of-the-art drug discovery pipelines (*3, 5*). Here, we introduce GENEVA (for <u>gen</u>etically diverse and <u>e</u>ndogenously controlled phenotypic <u>v</u>ariation <u>a</u>ssay), a framework which enables scalable molecular phenotyping of tumor models both in culture and in xenografted mice. GENEVA utilizes high-resolution molecular data from various cancer cell models to capture both inter- and intra-population heterogeneity in drug response. By studying drug effects across a range of genotypes within an internally controlled and highly quantitative setting, GENEVA uncovers common and therefore generalizable consequences of drug action across tumors.

At its core, GENEVA uses high-resolution transcriptomic measurements of natural phenotypic variations across panels of cancer cell models. In a GENEVA experiment, cells from tens of genetic backgrounds are combined into a single pool that is then grown in 3D cultures or xenografted in mice. The resulting mosaic tumor models serve as the unit of observation and are subjected to chemical perturbations *in vivo* or *in vitro*. The entire pool is then single-cell profiled, and the identity of every cell is determined based on its known reference genotype (**Fig S1**). This approach simultaneously captures macroscopic, cellular, and molecular phenotypes across all patient-derived lines, aiding in the understanding of the genetic and molecular factors of drug response and resistance.

Here, we have applied GENEVA to gain insights into the action of the recently approved Ras inhibitors approved in 2021, specifically the compounds targeting the oncogenic KRAS G12C mutation (*6, 7*). Although these compounds have recently invigorated the field of RAS targeting agents by their rapid development and approval into the clinic, several key challenges and gaps in our knowledge remain. Notably, the first approved KRAS G12C inhibitor Sotorasib has a median progression free survival (PFS) of only about 6 months, leaving much to be desired both in terms of PFS as well as resistance mechanisms which seem to occur in refractory patients (*8, 9*). By creating mosaic populations composed of KRAS variants (G12C and non-G12C) in culture and *in vivo* (both cell-line and patient-derived xenograft models), we revealed several novel insights into the biology of these Ras inhibitors and uncovered new molecular mechanisms cancer cells employ to achieve tolerance: (i) a previously unknown role for mitochondrial activity downstream of RAS inhibition in causing cell death, (ii) implication of multiple key pathways in drug tolerance, and (iii) identification of resistance mechanisms that emerge strongly when studied within the *in vivo* context. In addition to showcasing the power of GENEVA in augmenting our understanding of how chemistry perturbs biology, these findings lay the groundwork for advancing the emerging class of targeted RAS inhibitors and expanding combination therapies that address these diverse biological mechanisms.

## Results

### Multiplexed rich phenotyping to identify determinants of drug response and tolerance

We first evaluated GENEVA’s capacity to capture cellular and transcriptomic responses in diverse genetic mosaic populations. Initially, we pooled 11 cell lines, including ones with BRAF(V600E) and KRAS G12C mutations known for their sensitivity to Vemurafenib and ARS-1620, respectively (*10, 11*). Vemurafenib is an actively used inhibitor of BRAF(V600E) mutant melanomas in the clinic and ARS-1620 is one of the earliest developed inhibitors of KRAS.G12C developed by the Shokat group (*12, 13*). We then split this pool into three treatment conditions: DMSO, Vemurafenib, and ARS-1620 and used MULTIseq reagents to hash individual samples and combine them into a single scRNA-seq run (*14*). Next, we used SNP-based deconvolution and hash oligo demultiplexing to identify the genetic identity and the treatment condition for each profiled cell (**Fig 1A** and **Fig S1B-C**) (*15*). We readily noted several expected patterns, such as A375 cells (BRAF V600E mutant, red dashed outline in **Fig 1A**) being highly depleted in response to Vemurafenib (*16*). Similarly, for the ARS-1620 treated pool, we detected substantially fewer MIA PaCa-2 cells (KRAS G12C mutant) in the treated population (**Fig S1D**) (*17*). More importantly, we also observed clear patterns of intra-population heterogeneity in the sensitivity of MIA PaCa-2 cells to ARS-1620. As shown in **Fig S1D**, the distribution of ARS-1620 treated MIA PaCa-2 cells across the possible gene expression landscape significantly deviates from random chance (z-score of 6.65), highlighting strong heterogeneity in response.

**Figure 1.**
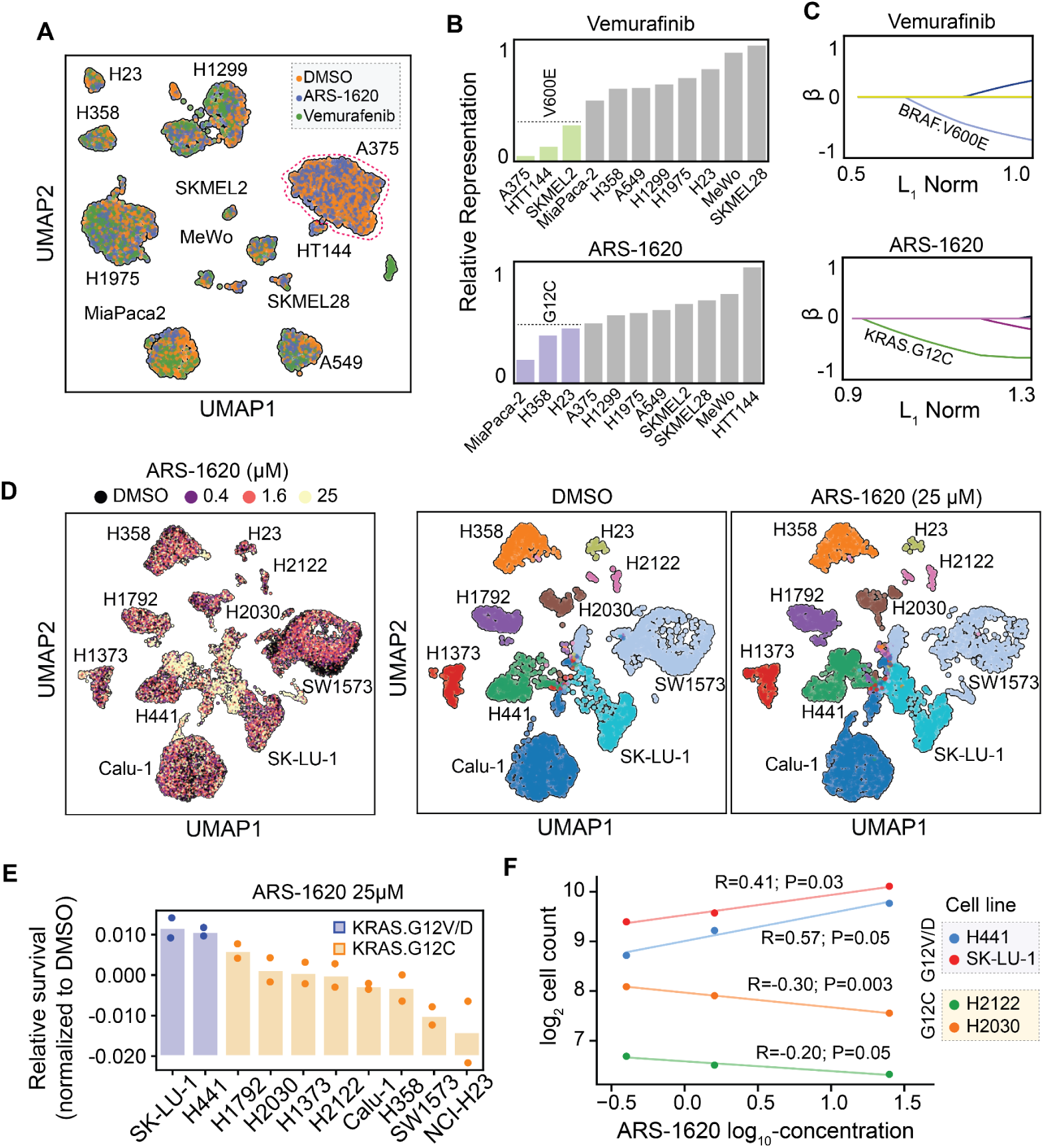
GENEVA allows for rich phenotypic profiling of cells in genetically diverse populations. **(A)** Our GENEVA pilot study with 11 human cell lines and three treatment conditions: DMSO, ARS-1620 (1 µM), and Vemurafenib (0.8 µM) treatment for six days. The UMAP projection summarizes the resulting single cell RNA-seq data. Cells are colored based on their drug treatment and the cell line identity of each cluster is labeled. The A375 cluster is marked by a dashed red circle. **(B)** The fraction of cells in each cell line in the treatment vs control populations were used to quantitatively measure the sensitivity of each line. Cell counts are from the genotyped scRNAseq data. Lines are sorted based on their relative sensitivity and G12C and V600E mutant lines are marked in ARS-1620 and Vemurafenib plots respectively. Relative representation was calculated by normalizing the cell counts by the cell line by the maximum number of cells in each respective drug condition, this ratio is shown. **(C)** Lasso regression with overlapping genetic mutations as covariates and GENEVA relative survival data as outcomes. Variables showing selection by the model are colored and labeled. KRAS G12C and BRAF V600E mutations are marked in each plot. **(D)** The summary of a G12C GENEVA pool composed of eight G12C cell lines along with 2 non-G12C lines (G12V and G12D) and three doses of ARS-1620 treated for six days. Shown are the UMAP projections for the DMSO and 25 μM ARS-1620 respectively. **(E)** Cell line representation at the highest dose relative to DMSO was used to measure relative drug sensitivity for each sample. Results are shown as a sorted bar graph where individual replicates are included as points with the bar showing the mean of the data (n=2). **(F)** Comparison of cell representation across doses for two G12C and two non-G12C lines. Lines are colored by identity and grouped into G12C and non-G12C lines with G12C lines showing negative slope across dose titrations of ARS-1620.

These observations show that both the number and state of cells in GENEVA effectively capture phenotypic traits such as sensitivity and response heterogeneity (**Fig S1E**). Cell counts were highly reproducible between biological replicates for all cell line-drug combinations. Additionally, we observed an expected correlation between relative cell counts and growth rates in the control sample, which led to our decision to balance future GENEVA pools, i.e. optimize the ratio at which the various lines are mixed based on cell proliferation rates, to ensure more equal representation of all lines at the time of data collection (**Fig S1F**).

We further analyzed the relationship between population representation and drug sensitivity by comparing cell lines under vehicle and drug treatments. As expected, BRAF V600E mutants were most sensitive to Vemurafinib and KRAS G12C mutants to ARS-1620 (**Fig 1B** and **Fig S1G**). More importantly, lasso regression analysis, with GENEVA-based relative sensitivities as response and whole-exome derived somatic mutations as covariates, correctly captured BRAF V600E and KRAS G12C as drivers of drug sensitivity to Vemurafinib and ARS-1620, respectively (**Fig 1C**).

Having observed clear inter- and intra-population variations in response to ARS-1620 among KRAS G12C lines, we sought to better understand the molecular basis of this heterogeneity. For this, we conducted a focused study using a pool of eight G12C and two non-G12C lung cancer cell lines. We treated these GENEVA pools with three concentrations of ARS-1620 for six days, performed single cell RNA sequencing, and analyzed the genotype of each cell as before (**Fig 1D** and **Fig S1H**).

Comparison of the two replicates across treatments and cell lines showed highly reproducible cell counts (R=0.9, **Fig S1I**). We also observed expected G12C-dependent changes in pool representation, with non-G12C lines becoming relatively more abundant at higher ARS-1620 concentrations (**Fig 1E**). Furthermore, akin to a dose response analysis, the inclusion of multiple concentrations of ARS-1620 in this GENEVA experiment allowed us to assess changes in pool representation as a function of drug concentration. For example as shown in **Fig 1F**, the relative pool representation of the non-G12C lines, H441 and SK-LU-1, increased across ARS-1620 concentrations; whereas H2122 and H2030, which are sensitive G12C lines, decreased. We calculated a sensitivity score for each line by measuring the slope in the log-log plots presented and observed a clear demarcation between G12C and non-G12C lines as well as a range of sensitivity among the KRAS G12C mutant backgrounds (**Fig S1J**).

### *In vivo* GENEVA for multiplexed high content phenotyping in xenograft models

The scalability of *in vivo* oncology xenograft models is particularly limited by the inability to effectively test multiple perturbations across many models simultaneously. This is due to technical variations between xenografted mice, which in turn necessitates large numbers of animal subjects to achieve sufficient power. By providing an endogenously controlled framework in which individual cells become the units of observation, GENEVA can minimize unwanted technical variations *in vivo* and reproducibly collect data across genetic backgrounds. We therefore set out to adapt GENEVA for *in vivo* studies using established patient and cell-line derived xenograft (PDX and CDX) models (**Fig S2A-B**). We initially created flank xenograft mosaic tumors comprising various KRAS G12* mutant cell lines across multiple cancer indications (**Fig S2A, S2C**). Following injection of the mosaic tumor, we treated the mice with either ARS-1620 (100 mg/kg) or the vehicle control for 10 days. Post-treatment, we resected the tumors, dissociated them into single-cells, enriched for cancer cells through a mouse cell removal step, and carried out single-cell RNA sequencing (**Fig 2A** and **Fig S2C**). Using relative cell counts, we then performed gross phenotyping of lines across KRAS G12* mutations. As expected, we observed significantly fewer cells from the KRAS G12C mutant lines after ARS-1620 treatment (**Fig 2B, Fig S2D**). More importantly, given the gene expression nature of our read-out, we also used annotated cell-cycle genes to examine the effect of ARS-1620 on cell cycle phase across mosaic xenografts (**Fig S2E**). Consistent with the known cytostatic effect of KRAS inhibitors (*18*), we found that in some cases (e.g., H2122), in addition to lower cell counts (**Fig S2F**), KRAS G12C inhibition also resulted in G1 arrest as measured by the proportion of cells in G1, G2M, and S (**Fig 2C**).

**Figure 2.**
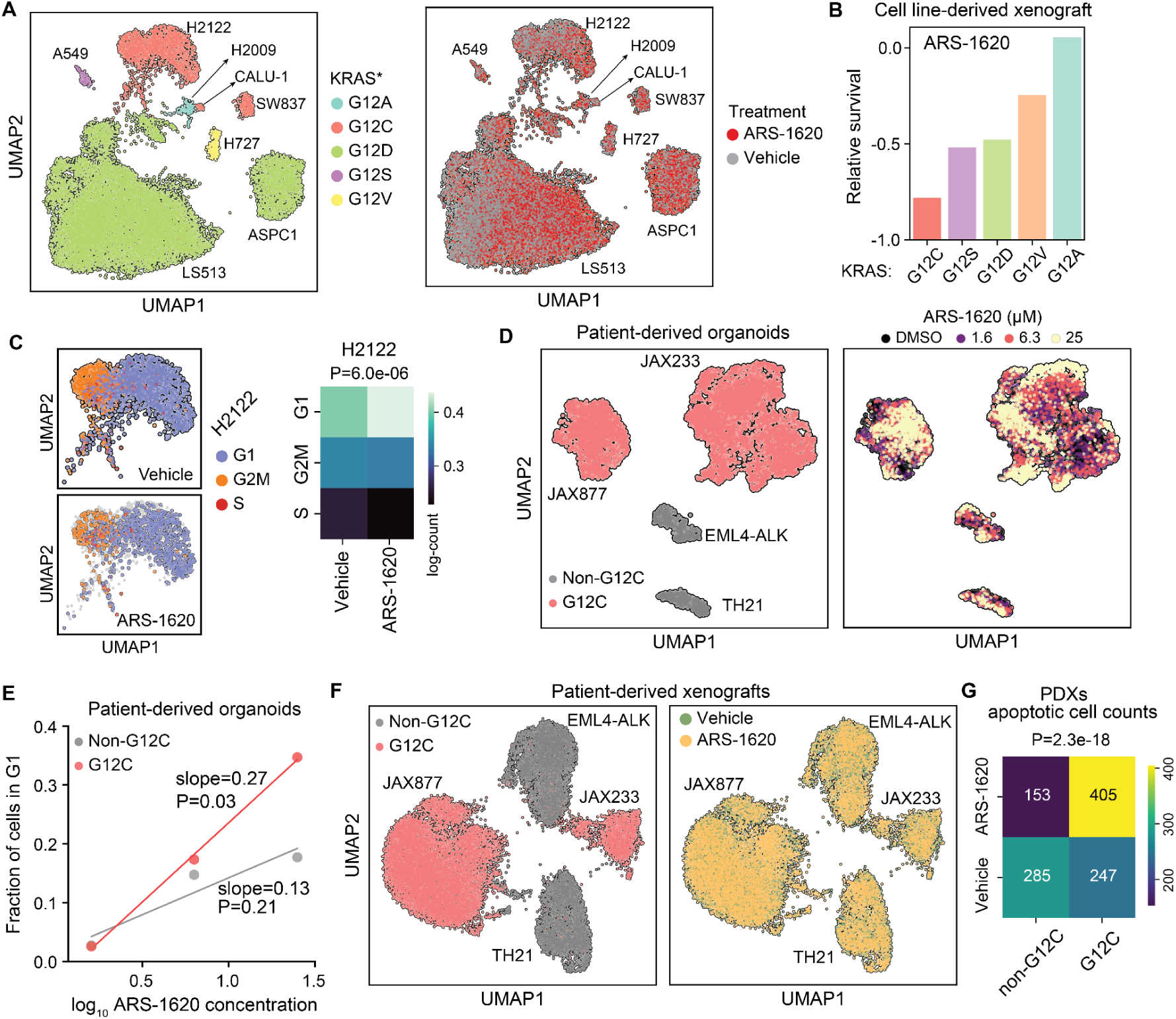
*In vivo* GENEVA applied to KRAS.G12 cell line- and patient-derived xenograft models. **(A)** NSG mice xenografted with mosaic cell-line models from a pool of KRAS G12* mutants were treated with ARS-1620 (100 mg/kg) or vehicle control for ten days after injection. Individual cells are colored by either KRAS G12* mutant status (left) or the treatment condition (right). **(B)** *In vivo* sensitivity phenotypes from CDX GENEVA mosaic tumors. Relative survival scores were calculated for each cell line by comparing their counts in drug and vehicle conditions and aggregated across different KRAS G12 mutations. **(C)** The UMAP projection of H2122 cells in the CDX GENEVA dataset colored by cell cycle phase (left). The number of cells in each phase was tabulated for ARS-1620 and vehicle-treated samples (right). *P*-value was calculated using a χ² test. **(D)** UMAP projection of mosaic PDO in an ARS-1620 GENEVA experiment. The cells are colored by G12C status (left) and ARS-1620 dose (right). Organoids were treated for 96 hours. **(E)** The relationship between fraction of cells in G1 and ARS-1620 concentration for G12C (red) and non-G12C PDOs. Shown are the slopes and their associated *p*-values. **(F)** UMAP projection of mosaic PDX in an ARS-1620 in-vivo GENEVA experiment where NSG mice were treated with ARS-1620 (100 mg/kg) for sixteen days. Cells are colored by G12C status (left) and ARS-1620 or vehicle treatment (right). **(G)** The number of cells in apoptotic cell state were counted for ARS-1620 and vehicle treatment samples across KRAS G12* mutation status. *P*-value was calculated using a χ² test.

Moving beyond cell lines, we expanded GENEVA to patient-derived models, both in 3D organoid (PDO) and in xenograft (PDX) forms, which are often considered a gold-standard in cancer studies (*19*) based on their higher fidelity in representing patient tumors (**Fig S2B**). For this purpose, we selected four lung cancer PDX models, two with G12C mutations (JAX233 and JAX877), one with an EML4-ALK fusion, and one with EGFR.L858R (TH21). We created GENEVA organoid pools using these models, treating them with varying ARS-1620 concentrations (**Fig 2D** and **Fig S2G**). In this case, as shown in **Fig 2E** (and **Fig S2H**), the cells from the G12C models showed a significantly higher dose-dependent G1 arrest when treated with ARS-1620 relative to non-G12C models.

Finally, we tested *in vivo* GENEVA on the four PDXs mentioned above in mosaic flank xenografts. Similar to the PDO experiment, mixed populations of cells were injected into mice, and once tumors were formed (48 hours), mice were treated with 100 mg/kg ARS-1620 or vehicle control (2 mice per condition) for 16 days. In this case as well, we observed robust representation of all four models within the mosaic tumors (**Fig 2F** and **Fig S2I**). It should be noted that unlike the *in vitro* cell line experiments, a significant change in the number of cells was not expected within the duration of these experiments. However, we observed a significant increase in the number of apoptotic cells among the G12C PDXs in the ARS-1620 treatment condition (**Fig 2G** and **Fig S2J**).

Taken together, our findings showcase GENEVA as a versatile platform, applicable to highly multiplexed mosaic pools both *in vivo* and *in vitro*. The resulting datasets allow for direct assessment of various phenotypic scores across different genetic backgrounds, treatments, and doses. However, absent from this picture is the utilization of the rich transcriptomic phenotypes and intra-population variations that were commonly observed in our data. In the following sections, we demonstrate that intra- and inter-population heterogeneity in drug response captured by GENEVA provides insight into molecular mechanisms of drug action and tolerance.

### KRAS G12C inhibition increases the activity of the mitochondrial electron transport chain

With access to multiple G12C-focused GENEVA datasets across various models, we sought to better understand the universal mechanisms of resistance to these therapies both *in vivo* and *in vitro.* First, we focused on data from our *in vitro* G12C focused pool with multiple doses of ARS-1620 (**Fig 1D**). At first glance, we made the interesting observation that KRAS G12C mutant cells that survived treatment with ARS-1620 had significantly fewer mitochondrial transcripts compared to their untreated counterparts, a signal that was overall consistent across all KRAS G12C lines. (**Fig 3A** and **Fig S3A**). Moving towards gene-level analysis, we performed differential gene expression analysis between treated and untreated cells across all KRAS G12C lines in order to reveal common effects of KRAS inhibitors. Consistent with our earlier observation, gene-set analysis of the resulting gene expression changes revealed that transcripts of mitochondrial-origin, but not genomic-encoded mitochondrial targeted transcripts, were significantly downregulated after ARS-1620 treatment (**Fig 3B**). To ask whether silencing mitochondrial genes resulted in an increase in survival upon ARS-1620 treatment, we also analyzed two previously published genome-scale ARS-1620 synthetic lethal CRISPR-interference screens (*20*). As shown in **Fig 3C** and **Fig S3B**, we observed that in H358 and MIA PaCa-2, which are both G12C lines, knockdown of mitoribosomal and mitochondria-associated genes (mitocarta) genes significantly increased survival (*17, 21*). To ensure that this effect is specific to ARS-1620, we analyzed several similarly generated synthetic lethal CRISPRi datasets (*22–24*). We also included a genetic screen for ATP levels as a positive control (*25*), and as expected, silencing mitoribosomal genes resulted in a significant reduction in ATP in this dataset (**Fig S3C**). Of the datasets we analyzed, none phenocopied the ARS-1620 screens (**Fig S3C)** highlighting the ARS-1620 specificity of mitochondrial involvement in sensitivity. Together, these results indicate that inhibiting mitochondrial function or reducing mitochondrial content results in increased tolerance to ARS-1620.

**Figure 3.**
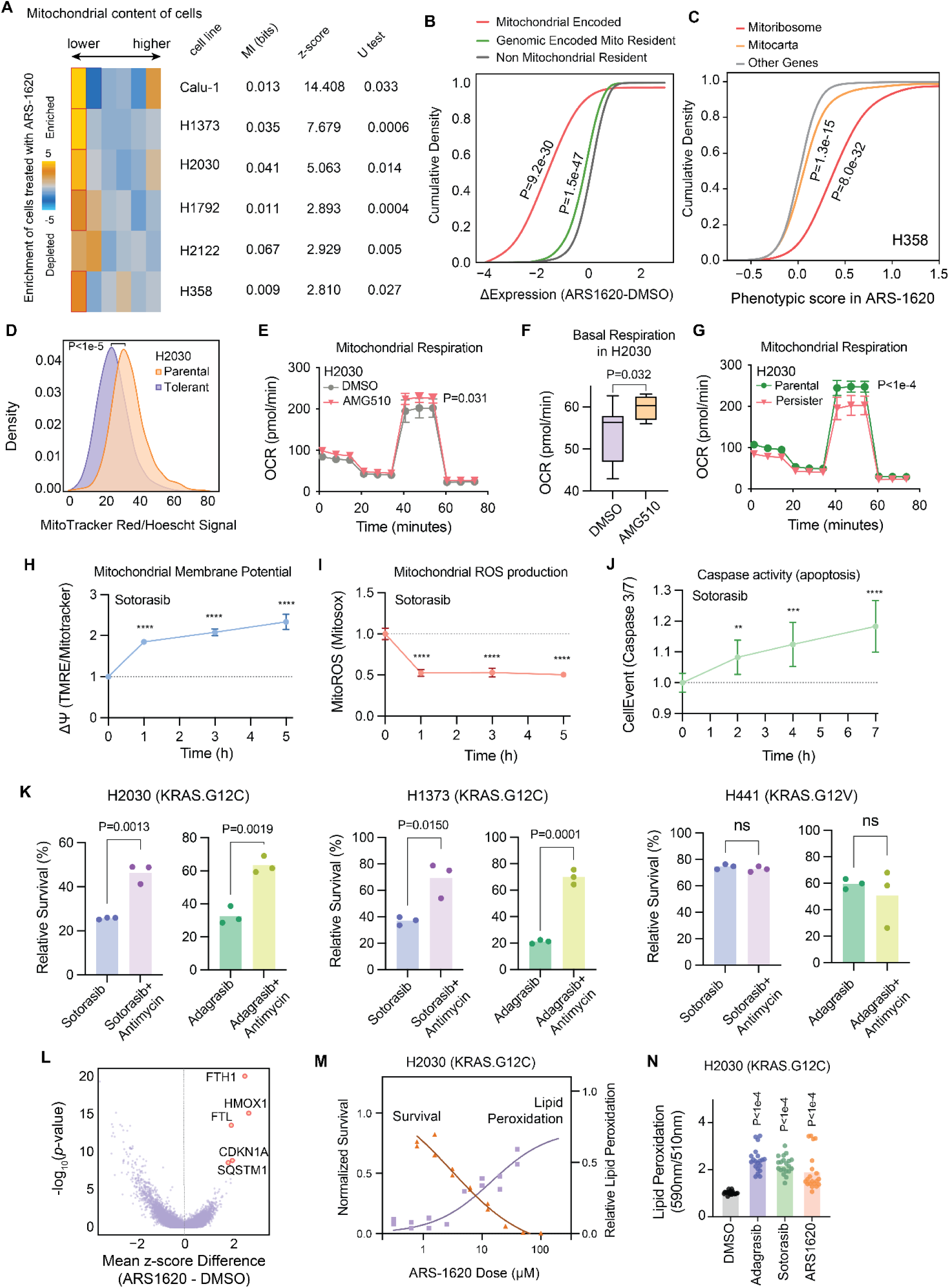
KRAS G12C inhibition leads to cell death via mitochondrial respiratory activity. **(A)** Cells surviving ARS-1620 treatment are enriched among those with lower mitochondrial content. For this analysis, cells were binned by expression of mitochondrial genes from low content (left) to high content (right). For each cell line in the GENEVA pool, the number of cells that were treated by ARS-1620 and reside in each bin was used to calculate an enrichment (gold) or depletion (blue) score. The significance of the pattern was measured using mutual information and its associated z-score as well as a two-sided Mann-Whitney U test. **(B)** Differential expression upon ARS-1620 treatment across mitochondrial encoded, genomically encoded but mitochondrial resident, and non-mitochondrial resident protein coding gene sets visualized as cumulative distributions. Lower differential expression indicates downregulation post-treatment. Statistics were calculated using a two-sample t-test for each geneset using the non-mitochondrial resident genes as the null distribution. **(C)** Genome-wide CRISPR-interference screening data from KRAS G12C mutant H358 line in a synthetic lethal screen with ARS-1620 grouped by MitoCarta, Mitoribosome, and all other genes. Increase in phenotypic score indicates a buffering or rescuing effect on cell survival in a drug treatment context. Data from Lou *et al* (*20*). **(D)** H2030 persistor and parental cell lines profiled for mitochondrial content by MitoTracker Red. P calculated using two-tailed t-test. **(E)** Oxygen consumption rate (OCR) in H2030 measurements after acute treatment with Sotorasib for 2h at 500 nM using a seahorse assay. *P* values were calculated by two-tailed t-test (n=10). Shown is the mean and standard deviation. **(F)** Basal respiration in H2030 after acute treatment with Sotorasib, data from seahorse assay. *P* values were calculated by two-tailed t-test (n=10). Shown is the mean and standard error of the mean. **(G)** OCR measurements after acute treatment with Sotorasib for 2h at 500 nM with a seahorse assay in parental (green) and persistor (pink) lines. *P* values were calculated by two-tailed t-test (n=10). Shown is the mean and standard deviation. **(H)** Mitochondrial membrane potential measured by TMRE and MitoTracker Red ratio in flow cytometry assays after induction with Sotorasib (500nM) for 0-5 hours. **(I)** Mitochondrial ROS formation measured by Mitosox signal in flow cytometry assays after induction with Sotorasib (500 nM) for 0-5 hours. **(J)** Caspase 3/7 cleavage measured by CellEvent Caspase 3/7 signal in flow cytometry assays after induction with Sotorasib (500 nM) for 0-7 hours. For H-J, for each timepoint, thousands of cells measured from two wells were sampled and bootstrapped with equivalent numbers of cells from each well without replacement (bin size = 0.02) until all cells were fully binned; each randomly sampled bin is plotted as a replicate. Shown is the mean +/- standard deviation. **(K)** Cell survival proportions with Sotorasib and Adagrasib combination dosing with Antimycin A in two KRAS G12C cell lines (H2030, H1373) and one non-G12C cell line (H441). *P* values were calculated by two-tailed t-test. Barplots represent the mean of the data (n=3). **(L)** Volcano plot visualizing differentially expressed genes across KRAS G12C cell lines in GENEVA pools after drug treatment with ARS-1620. P-values are calculated by two-sample t-test scoring independent cell lines as biological replicates. **(M)** Relative cell survival and lipid peroxidation dose-response to ARS-1620 after three days of treatment as measured by CTG and BODIPY 581/591 C11. The left hand Y-axis for survival, subsequently plotted as orange data points, and the right hand Y-axis for lipid peroxidation, subsequently plotted as purple data points. **(N)** Lipid peroxidation levels, measured by BODIPY 581/591 C11, after treatment with three KRAS G12C inhibitors compared to a vehicle. *P* values were calculated by two-tailed t-test (n=20).

To further explore this mitochondrial phenotype, we generated an ARS-1620 persister derivative of the H2030 cell line by treating this KRAS G12C line with ARS-1620 (10 µM) for 34 days. Consistent with our observations from GENEVA, measurement of mitochondrial content using Mitotracker-DeepRed showed a significant decrease in the mitochondrial content of the H2030 ARS-persister line compared to the drug-naïve parental cell line (**Fig 3D**). Given this observation, we next sought to better capture the mechanistic effects of KRAS G12C inhibitors on mitochondrial dynamics. To ensure the KRAS specificity of this signal beyond ARS-1620, we turned to the more recent clinically approved KRAS G12C inhibitor Sotorasib, which has a sub-micromolar IC50. Using a Seahorse Respirometer (*26*), we measured oxygen consumption dynamics with and without Sotorasib treatment in a two-hour post-treatment window. Strikingly, we observed that Sotorasib induced a rapid and significant increase in basal respiration (P=0.03, **Fig 3E**). This effect is attributable to the increase in spare respiratory capacity at the site of the electron transport chain, with no significant change in non-mitochondrial derived respiration or ATP production (**Fig 3F** and **Fig S3D-E**). Interestingly, the increase in spare respiratory capacity attributable to Sotorasib was absent in H2030 persister cells, which further highlights the epistatic interaction between mitochondrial function and KRAS inhibition (**Fig S3F**) and points to a potential adaptive mechanism present in persister cells. Persister cells also displayed a higher baseline spare respiratory capacity compared to parental cells suggesting a further compensatory mechanism of increased mitochondrial activity to balance fewer mitochondria. This marked increase in mitochondrial respiration was unexpected as most inhibitors that perturb the electron transport chain cause a decrease in mitochondrial respiration (*27*). Additionally, this acute increase in respiration was lost in treatment persister cells suggesting that this mechanism is selected against during drug treatment. We further characterized this mitochondrial hyperactivity phenotype by conducting an acute time course study measuring mitochondrial membrane potential, mitochondrial reactive oxygen species (mitoROS), and caspase cleavage in non-persister cells using flow cytometry sensors. The initial cellular response to KRAS G12C inhibition was marked by an increase in mitochondrial membrane potential (TMRE) at one hour after treatment, consistent with the observed acute upregulation in mitochondrial oxygen consumption (**Fig 3H**). At the same time, mitochondrial ROS production (MitoSox) decreased at one hour after treatment while Caspase 3/7 cleavage steadily increased between 0-7 hours post treatment (**Fig 3I, 3J**). While caspase 3/7 cleavage has been shown to result from KRAS G12C inhibition, our data show that strong mitochondrial hyperactivity precedes that process suggesting that this phenotype precedes traditional mechanisms of cell death.

Given the impact of KRAS G12Ci on induction of elevated and aberrant mitochondrial activity in the acute phase of response to drug treatment, we hypothesized that Sotorasib may be acting via the mitochondrial electron transport chain (ETC) to induce cell death. To investigate this possibility we utilized Antimycin A1, a potent inhibitor of Complex III, to determine if selective inhibition of the rate of electron flux through the ETC could rescue lethality of Sotorasib in KRAS G12C mutant cell lines. We combined treatment of the two KRAS G12C inhibitors Sotorasib (Sotorasib) and Adagrasib (Adagrasib) with Antimycin A1 in CellTiter-Glo (CTG) survival assays. The results demonstrated that inhibition of ETC activity using Antimycin A1 attenuates lethality of the KRAS G12C inhibitors (**Fig 3K**). In H2030 and H1373 cells, two lung cancer cell line models with KRAS G12C mutations, we observed increased cell survival after treatment of KRAS G12C inhibitors in combination with Antimycin A1. In contrast, H441, a KRAS G12V lung cell line, did not show a significant change in cell survival. Taken together, the data demonstrates that KRAS G12C inhibitors induce cell death via a previously unappreciated mechanism of increased mitochondrial activity and that this mechanism is specific to KRAS G12C mutants, indicating that this is an on-target effect of KRAS G12C inhibitors.

A secondary signal derived from our focused KRAS G12C GENEVA pool suggested that KRAS G12C inhibitors could also be inducing ferroptosis as a mechanism of action. Across KRAS G12C lines surviving drug treatment, we found a striking upregulation of anti-ferroptotic genes including the two components of the ferritin complex FTH1 and FTL (**Fig 3L**). We further investigated this signal by measuring the amount of lipid peroxidation induced by ARS-1620 using the flow cytometry live-cell lipid peroxidation sensor BODIPY C11. We observed that ARS-1620 induced lipid peroxidation in a dose-dependent manner (**Fig S3G**) and that cell survival decreased as the amount of lipid peroxidation increased (**Fig 3M**). A comparison of the dynamics of these two phenotypes, survival and lipid peroxidation, between ARS-1620 and the well known ferroptosis inducer Erastin showed similar relationships between decreased survival and increased lipid peroxidation in a dose-dependent manner for both molecules (**Fig S3H**). To characterize the generalizability of this phenotype to multiple KRAS G12C inhibitors we performed similar lipid peroxidation measurements (*28*) in response to Adagrasib, Sotorasib, and ARS-1620 in KRAS G12C mutant and non-G12C mutant cell lines. We found that all three compounds significantly increased lipid peroxidation in a KRAS G12C mutant line H2030 (**Fig 3N**) while this effect was significantly decreased in a non-KRAS G12C model H441 (**Fig S3I**). To assess if induction of ferroptosis was a significant contributor to cell death in response to on-target effects of KRAS G12C inhibitors, we sought to rescue cell lethality of Sotorasib and Adagrasib using the anti-ferroptotic agent Ferrostatin-1 (*29*). However, unlike Antimycin, Ferrostatin-1 was unable to rescue the lethality of KRAS G12C inhibitors indicating that although KRAS G12C inhibitors were able to induce ferroptotic markers, this was not a primary mechanism of action of how the molecules impacted cell survival (**Fig S3J**).

These two examples illustrate how GENEVA is able to uncover primary and secondary mechanisms of drug action. GENEVA allows for stratification on both phenotypic and molecular levels, which allows us to link specific signals to key cellular pathways and processes: (i) genotype stratification between sensitive (KRAS G12C) and insensitive (non-KRAS G12C) lines, (ii) intra-population heterogeneity analysis by single-cell methods to profile drug-tolerant cells within populations, and (iii) cross-model generalizability by examination of shared signals between all sensitive genotypes. By combining analysis on multiple levels from a single GENEVA experiment we were able to find previously unknown mechanisms for highly characterized molecules demonstrating the utility of GENEVA in drug discovery contexts.

### Discovering and targeting the mechanisms of tolerance to KRAS G12C inhibition *in vivo* and *in vitro*

As we showed in **Fig S1D**, there is apparent heterogeneity in drug sensitivity within KRAS G12C models. We posited that comparing these surviving sub-populations across all sensitive lines reveals general mechanisms of resistance to KRAS G12C inhibition. We therefore performed a differential expression analysis focused on genes that are differentially expressed in ARS-1620 treated cells in a KRAS G12C-dependent manner (**Fig 4A** and **Fig S4A**). In this analysis, we observed higher expression of several druggable protein-coding genes that point to their higher activity conferring resistance in KRAS G12C mutant cells including: JAK1, RPTOR, PARP14, GPX4, AURKA, EHMT2, and IDH2. Several of these pathways, such as mTOR and AURKA, have already been implicated in drug resistance.To test this possibility, we used a panel of compounds targeting these respective pathways, namely Ruxolitinib (JAK), INK128 (mTOR), RBN012759 (PARP), RSL3/Erastin/Altretamine (Ferroptosis), Alisertib (AURKA), BIX01294 (Histone methyl-transferases), and Enasidenib (IDH2) to assess potential synergies with KRAS inhibition (*30, 31*). As shown in **Fig 4B**, in the case of RBN012759, Axitinib, Enasidenib, and INK128, we observed a significant synergy as measured by Bliss scores across three KRAS G12C inhibitors (*32*). In contrast, consistent with our observations with Ferrostatin-1, inducers of ferroptotic cell death did not show a significantly synergistic interaction with Ras inhibition (**Fig S4B**). For INK128, which showed the strongest synergy, we also performed an *in vivo* combination therapy with subcutaneous H2030 tumors in xenografted mice. Once tumors were formed (21 days post-injection), the mice were randomly divided into four arms, and were treated with ARS-1620 (100 mg/kg) and INK128 (0.3 mg/kg), both as monotherapies and in combination. While these drugs were effective in reducing tumor growth in isolation, consistent with our observations *in vitro*, their combination showed a substantial and significant increase in efficacy (**Fig 4C** and **Fig S4C**). Taken together, our results indicate that the cancer cell heterogeneity revealed by GENEVA can be harnessed to identify generalizable mechanisms of drug resistance and design rational combination therapies for improving the efficacy of targeted therapeutics.

**Figure 4.**
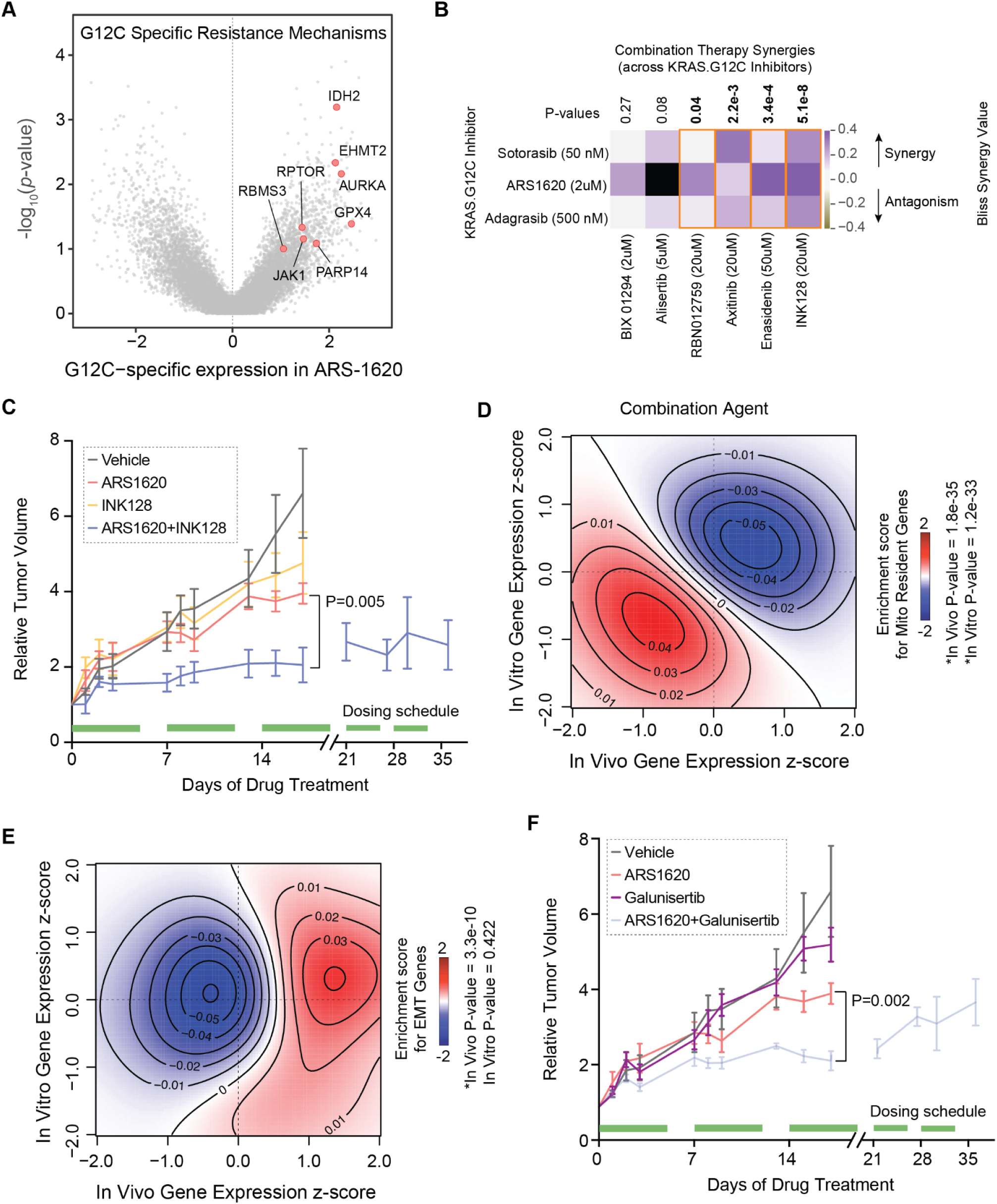
Targeting pathways associated with increased tolerance to KRAS G12C inhibition leads to more efficacious combination therapies. **(A)** Volcano plot of G12C showing KRAS G12C specific gene expression upregulation. G12C specific expression was taken by first calculating differential expression between DMSO and ARS-1620 treatment for each cell line and then taking the mean log2 fold-change difference between cell lines with G12C and non-G12C mutations. *P-values* were calculated using two-tailed t-tests between G12C and non-G12C groups. Druggable proteins significantly upregulated in a G12C specific manner are highlighted in red. **(B)** Combination therapy testing with three KRAS G12C inhibitors and compounds targeting G12C-specific resistance mechanisms identified in Fig 4A was performed in a three-day CTG survival assay in H2030. Data is shown with Bliss synergy values colored as a heatmap with purple indicating Bliss drug synergy and grey indicating Bliss antagonism, with an orange outline indicating p-values less than 0.05. *P-values* were calculated by two-tailed t-test of replicates specific to a dual-agent combination compared against a null distribution estimated from no drug biological replicates (n=4). **(C)** Combination therapy *in vivo* study performed in NSG mice with treatment duration between 17-35 days. ARS-1620 (100 mg/kg) was co-administered with the mTOR inhibitor INK128 (0.3mg/kg). ARS-1620 and INK128 were dosed singly and in combination daily during indicated dosing schedule intervals (green) by IP injection with n=5 NSG mice per group. Shown are mean +/- standard deviation. **(D)** Two-dimensional density plot showing concordance of gene expression downregulation between *in vivo* and *in vitro* mosaic pools. Red indicates mitochondrially encoded gene expression enrichment; blue indicates depletion. Expression data was transformed to differential expression z-scores by comparison of ARS-1620 to DMSO conditions for respective *in vitro* and *in vivo* GENEVA pools. *P-values* were calculated by Mann Whitney U-test of z-scores from genes within the geneset shown and all other genes. *P-values* indicate the significance of downregulation in each model system (vitro/vivo). **(E)** Two-dimensional density plot showing model specific downregulation of gene expression between *in vivo* and *in vitro* mosaic pools. Red indicates EMT gene-set enrichment; blue indicates depletion. Statistics were calculated as described in Fig 4D. **(F)** Combination therapy *in vivo* study with TGFB/EMT inhibitor Galunisertib (75 mg/kg) and ARS-1620 (100mg/kg). ARS-1620 and Galunisertib were dosed singly and in combination daily during indicated dosing schedule intervals (green) by IP injection with n=5 NSG mice per group. Shown are mean +/- standard deviation. Control and ARS-1620 mice data is shared between Fig4C and Fig4F. Study design and statistics were as described in Fig 4C.

As we discussed earlier, GENEVA allows for rich profiling of drug response both *in vivo* and *in vitro*. However, we had not addressed the extent to which these two modalities may differ and whether *in vivo* GENEVA, which better captures the cellular context of a tumor, provides greater insights into mechanisms of drug action and resistance. To tackle this, we performed a comparison of G12C-associated gene expression changes in response to ARS-1620 both *in vivo* and *in vitro*. We observed an overall agreement between these two modalities. For example, mito-resident protein coding genes, which as we showed in the previous section are significantly down-regulated in ARS-1620 tolerant cells, showed a similar pattern both *in vivo* and *in vitro* (**Fig 4D**). However, we made the intriguing observation that genes associated with epithelial-to-mesenchymal transition (EMT) were significantly enriched among those up-regulated in ARS-1620 tolerant cells *in vivo* but not *in vitro* (**Fig 4E**). This observation indicated that cells undergoing EMT may be more resistant to KRAS G12C inhibition *in vivo*, an effect that is not as strongly and universally observed *in vitro*. Consistently, we also noted a similar up-regulation of EMT genes in the two ARS-1620-treated KRAS G12C PDX models presented in **Fig 2F** (**Fig S4D**). We posited that this phenotypic divergence between the two modalities may result from the entanglement between EMT and hypoxia, which is common in tumors (*33*). To test the potential role of EMT in response to KRAS G12C inhibition, we used ARS-1620+Galunisertib (TGF-β receptor inhibitor (*34*)) combination therapy to simultaneously target the EMT and Ras pathways. As shown in **Fig 4F** and **Fig S4E**, we observed a significant and synergistic reduction in tumor growth with this combination therapy in xenografted mice treated with 75 mg/kg of Galunisertib. Together, our results establish that (i) EMT plays a role in resistance to Ras inhibition, and (ii) *in vivo* GENEVA, which captures drug response in the context of the tumor, reveals additional pathways that may go unnoticed *in vitro*.

We further sought to interrogate the value of doing multiple models and using different biological models as biological replicates in our statistical tests for geneset and pathway discovery. We started by downsampling from the full number of KRAS G12C models in our mosaic from eight cell lines to two cell lines and asked if we reduced the group size if we would see a decrease in the number of differentially significant pathways as scored by GSEA. As we decreased the number of lines sampled as biological replicates we saw a concomitant decrease in the number of overlapping significant genesets, indicating that inclusion of multiple models in this type of framework allows for detection of shared significant signals despite underlying differences in gene expression changes **(Fig S3K, S3E).** As a specific illustration of the utility of inclusion of multiple models in discovering significant genesets more quickly, we performed a similar analysis but calculated the change in significance as measured by the p-value of each gene within genesets as scored from differential expression scoring. We found that for the two genesets which led us to the mitochondrial and mTOR hypotheses (Mito Encoded, Ribosomal genes) the significance increased as the size of the biological model set increased. The null dataset taken as a random sampling of genesets did not increase dramatically in significance **(Fig S3F).**

### Leveraging GENEVA for rich profiling of combination therapies

As shown above, single cell RNA sequencing provides a rich phenotyping strategy that goes substantially beyond the traditional cell-count based measures of drug sensitivity. The ability of GENEVA, especially in its *in vivo* modality, to successfully scale this type of molecular phenotyping prompted us to expand our approach beyond monotherapies. We asked whether GENEVA could be similarly leveraged to study *in vivo* drug synergies of combination therapies. To test this possibility, we created mosaic tumors composed of three KRAS G12C lines (CALU-1, H2030, and H1373) and a G12D line (LS513) as reference (**Fig S5A**). We treated mosaic tumor bearing mice with Antimycin, Galunisertib, and INK-128, both individually and in combination with ARS-1620. We then used *in vivo* GENEVA to create a large-scale map of phenotypic and molecular responses to these combination therapies (**Fig 5A**). We used the resulting dataset to study drug synergies at the molecular level. For example, we used our single cell RNA sequencing data to measure Bliss drug synergies in causing G1 arrest. As shown in **Fig 5B**, consistent with our observations in the previous sections, we observed a positive Bliss synergy between ARS-1620+Galunisertib and ARS-1620+INK-128 across the KRAS G12C models. In contrast, and as expected, Antimycin showed a directionally opposite synergy score in KRAS G12C cells, indicating rescue of G1 arrest when used in combination with ARS-1620. In order to identify the genes and pathways that contribute to these synergies, we used our scRNAseq expression data to fit linear models of gene expression based on the cell lines and drug treatment within the dataset to obtain gene level coefficients. In the same way that we previously modeled drug synergy on phenotype by a comparison of single-agent additive effects to dual-agent observed effects, we sought to find genes that fell outside of simple additive models. We fit a baseline model taken from data from single agent treatments in GENEVA pools and then fit a second model taken from data from single agent and dual agent treatments. For each gene we then calculated gene level coefficients from the single-agent model and compared these values to the coefficients from the dual-agent model derived from experimental dual-agent data. To identify genes that fall outside of an additive model, we ranked accordingly by their difference between single-agent and dual-agent models. Consistent with our earlier observations regarding the contribution of mitochondrial activity to the therapy resistance phenotype, here as well, mitochondrial genes were negatively downregulated in the dual-agent experimental conditions ARS-1620+Galunisertib and ARS-1620+INK-128 (**Fig 5C** and **Fig S5B**). Taken together, GENEVA not only allows for multiplexed phenotypic profiling of drug combinations *in vivo* but also allows for gene level understanding of which genes drive drug synergy.

**Figure 5.**
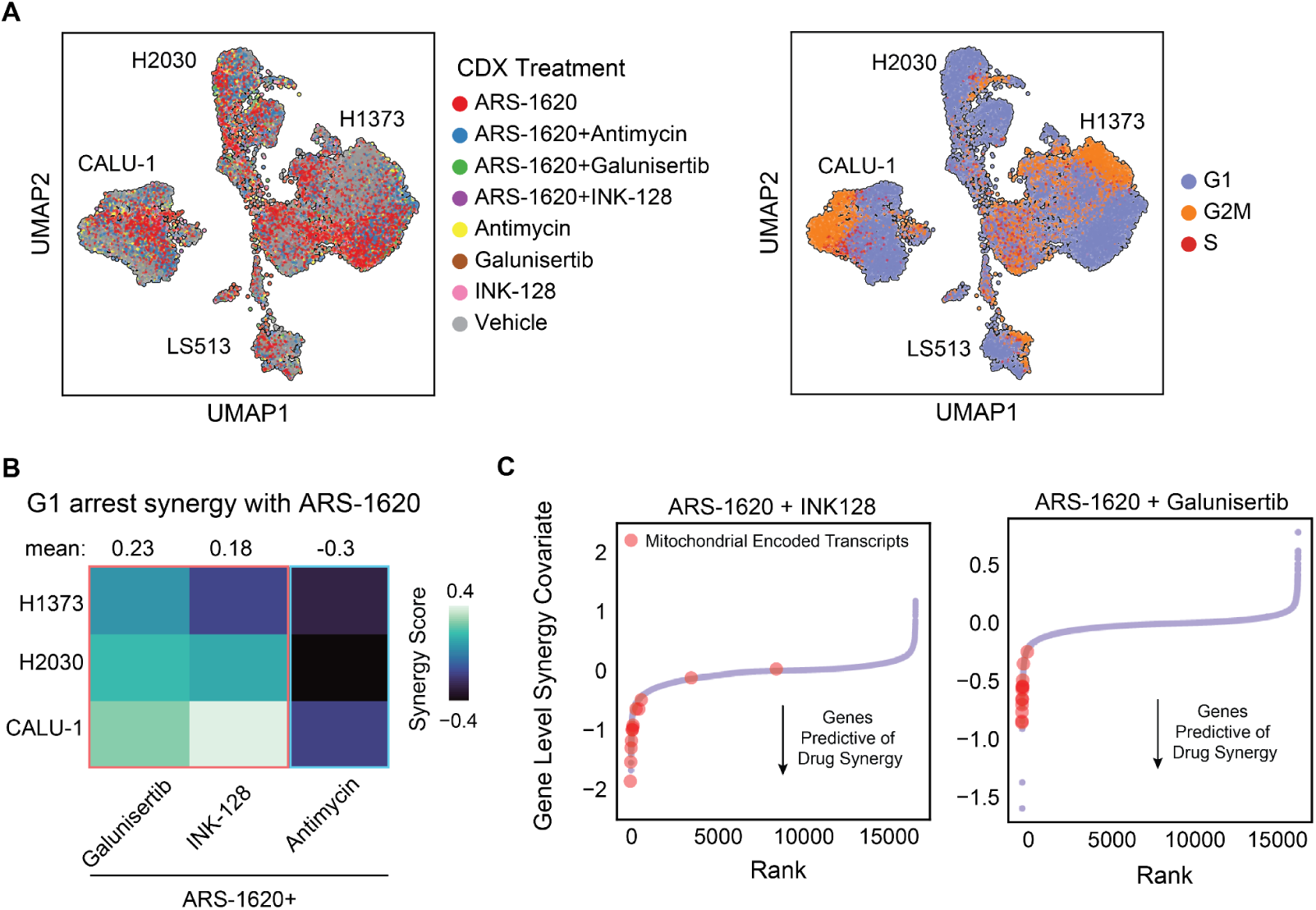
GENEVA enables multiplexed combination therapy screening *in vivo*. **(A)** Cell lines xenografted as mosaic tumors were treated with drugs in single or combination dosing (left). Mosaic tumors were grown for ten days before daily treatment with drug combinations for seven consecutive days by IP injection. ARS-1620 (100 mg/kg), Galunisertib (75 mg/kg), INK128 (0.3 mg/kg) and Antimycin A1 (0.5 mg/kg) were administered singly or in combination at these concentrations. Clusters are labeled by the cell line of origin and colored by the drug treatment condition. Cells are colored by their cell cycle state and identity (right). **(B)** Bliss synergy values calculated on G1 arrest scores for KRAS G12C cell lines from GENEVA after combination dosing with ARS-1620 and Galunisertib, INK-128, or Antimycin. Heatmap colors reflect the relative level of synergy for the two agents with lighter colors indicating synergistic suppression of G1 and darker colors indicating antagonistic effect on G1 phase. **(C)** Identification of genes which deviate from additive synergy models in terms of expression change in response to drug treatment. Data from single-agent and dual-agent GENEVA studies were used to build linear models for calculation of gene-level coefficients. The difference between single-agent and dual-agent model coefficients for each gene were calculated and rank-ordered. Z-scores of the value of the coefficient difference are plotted on the y-axis and the rank of the genes from lowest to highest plotted on the x-axis. This analysis was performed for both ARS-1620+INK128 and ARS-1620+Galunisertib. Mitochondrial genes are colored in red dots on both plots.

## Discussion

In this study, we introduced GENEVA, a framework that enables scalable molecular phenotyping of tumor models both *in vivo* and *in vitro*. We showed that GENEVA effectively reveals the molecular basis of inter- and intra-population heterogeneity in drug response by leveraging high-resolution measurements of natural phenotypic variations across a large panel of cancer cell lines and PDXs. While other methods have been developed that take advantage of multiplexed scRNAseq across models, drugs, and genetic manipulations, we have extended multiplexed scRNAseq to a scalable *in vivo* platform that can also utilize challenging model systems from patients (*35–37*). This extension, however, required multiple rounds of optimization and pool balancing to ensure sufficient representation of cells from each genetic background at the time of tumor harvesting, with the key insight for proper pool balancing relying on heuristics that inversely seed cells for accurate representation at the time of harvest. Data from cancer cells *in vivo* better captures the tumor microenvironment context in patients and the resulting is far more likely to generalize to human patients (*36, 38*). As we showed, there are key genes and pathways that emerge only when studied *in vivo*. The activation of the Ras signaling pathway is a hallmark of many cancers. Ras itself, which is recurrently mutated across many cancer types, is viewed as an Achilles heel of cancer cells. Long viewed as being undruggable, targeted therapies that bind and inactivate mutant forms of Ras are successfully entering the market after the key discovery of covalent inhibitors for KRAS G12C mutant cancers (*12, 39*). At the onset of our study, inhibitors of the KRAS G12C variant were making their way through clinical trials and given our longstanding interest in this pathway, we selected them for our first application of GENEVA.

We showed that a previously unappreciated mechanism of action of KRAS G12C inhibitors involved perturbation of mitochondrial respiration. Using a combination of mechanistic studies and genome-wide screening data we have shown that this observation can be causal in how KRAS G12C inhibitors successfully suppress cell growth. Based on our findings, KRAS G12C persister cells have less mitochondrial content (**Fig S5C**) and decreased mitochondrial respiration (**Fig 3G**). At an acute timepoint, we observed increased mitochondrial respiration in response to KRAS G12Ci. By combining treatment with an inhibitor of the mitochondrial electron transport chain (Antimycin), we were able to rescue lethality of KRAS G12Ci compounds demonstrating that this phenotype was key to cell death caused by KRAS G12C inhibitors. Furthermore, mining of synthetic lethal genome wide screen data demonstrated that negatively impacting mitochondrial biogenesis by knockdown of mitochondrial ribosomal genes rescues lethality to KRAS G12Ci (**Fig 3C**). Within the KRAS G12C space, studies examining the dynamics of KRAS G12C inhibition using small molecule targeted therapies have focused on upstream RTK reactivation or downstream rebound mechanisms (*40*),(*41*),(*42*); we propose that investigation of mitochondrial dynamics are another key area of focus in developing understanding of the mechanisms of action of these compounds and KRAS biology.

We also identified pathways of drug resistance (e.g. mTOR and EMT) that informed new combination therapies; two of which we validated *in vivo* using combination studies with ARS-1620. mTOR inhibition has been proposed as a key synergistic avenue for increasing response in patients in combination with KRAS G12C inhibitors and as a result several pharmaceutical companies are pursuing combination studies in clinical trials with KRAS G12Ci. Our platform has discovered and confirmed that this signal is robust in multiplexed GENEVA models. Recent data collected from patients supports our findings in regards to EMT for combination therapy as well. For example, in a study presenting different types of genetic or non-genetic alterations in patients refractory to Adagrasib, several patients displayed cellular transformation in refractory tumor biopsies accompanied by a lack of any novel genetic alterations (*43*). Notably, histology of these refractory tumors appeared to show a decrease in the number of tight and discrete epithelial foci which is also a hallmark phenotype of EMT. Studies using a combination of refractory cells derived from *in vitro* models and patient data also point to a cell morphology change after resistance to KRAS G12Ci treatment (*44*). Taken together with our finding that the upregulation of EMT was observed in KRAS G12C mutant cells surviving targeted therapy, there is an emerging hypothesis that cellular transformation is an important aspect of adaptation to KRAS G12Ci. We propose that EMT could be a driver of this resistance and have shown that inhibition of EMT synergistically improves outcomes *in vivo*.

Natural phenotypic variations in genetically diverse populations are a cornerstone of modern genetics (*45*). By scaling pharmacogenomic data collection across many models in an endogenously controlled and quantitative setting, GENEVA allows for a systematic search for genetic and molecular determinants of cellular response. As the number of genetic backgrounds in our mosaic tumors grow, tools developed for GWAS or QTL studies become readily applicable to data collected using GENEVA. More importantly, by collecting data at single cell resolution, GENEVA incorporates natural phenotypic variation among the population of cells within each population background. This allows us to derive insights from these datasets without the onerous sample size requirements that GWAS studies often contend with (*46*).

An important limitation of *in vivo* GENEVA is that, in its present iteration, the mosaic tumors are grown in immunocompromised mice. Therefore non-cell autonomous mechanisms of drug response that involve components of the adaptive immunity fall within the blindspot of this approach. Nevertheless, the vast majority of targeted therapies remain in the cell autonomous sphere with KRAS G12C mutated tumors being a notable example. There is also the remote possibility that cancer cells from different backgrounds may influence one another. In our experience, cancer cells in the pool grow and respond similar to their behavior in isolation. Furthermore, as the diversity of cells within the pool grows, the likelihood of a specific pair of cells coming to contact diminishes rapidly. Additionally, although we have demonstrated with multiple inhibitors and at multiple doses that GENEVA can be informative as to fitting models for detection of features that predict sensitivity, we believe it is a data-rich complement to many traditional methods such as dose-response etc.

Taken together, we believe that GENEVA has the potential to help us rethink how drugs are evaluated or even discovered. Through scalable molecular measurements across many diverse models and within their *in vivo* context, GENEVA allows us to capture heterogeneity in drug response much earlier in the drug discovery process. Moving beyond simplified biochemical or cell-based assays that are traditionally used for drug discovery provides an opportunity to identify drugs not solely based on their depth of target engagement, but also their breadth of activity across many models. Prioritizing this step ensures that the eventual drugs that make their way to the market are more likely to generalize across patients. In the same vein, rich phenotypic data from many tumor models allows us to better identify patients that are most likely to respond to each therapy, which in turn leads to more informed selection of patients in subsequent clinical trials. In our view, this is a crucial step towards realizing the promise of precision medicine.

## Acknowledgements

We thank the Laboratory Animal Resource Center (LARC) at UCSF. HG is an Era of Hope Scholar (W81XWH-2210121). Sequencing was performed on a HiSeq 4000 at the UCSF Center for Advanced Technologies, supported by UCSF PBBR, RRP IMIA, and NIH 1S10OD028511-01 grants.

We gratefully acknowledge the financial support provided by the National Science Foundation Graduate Research Fellowship Program (NSF GRFP) for JY. We thank Demian Sainz, Mark Ansel, and Anita Sil of the UCSF BMS program for their support for JY.

We thank Dr. Trever Bivona, Dr. Franzi Haderk, and Victor Olivas for their generous gift of the EML4-ALK mutant PDX model for this study.

KMS thanks the Howard Hughes Medical Institute, the Samuel Waxman Cancer Research Foundation, and NIH grants 1R01CA221969 & 5R01CA244550. KL thanks NIH grant F30CA239476.

Research in the lab of Dr. Hani Goodarzi is supported by the Arc Institute.

## Materials and Methods

### Organoid culture

PDX tumors were thawed from snap frozen vials in 10% DMSO, 90% FBS and digested with fine dicing by razor blade and digestion for 1 hour at 37C with 1mg/mL Liberase TM. Cell suspensions were strained in 100 micron filter filters and resulting cell suspension was seeded in 100% Matrigel at 37C on a dry sterile tissue culture plate. After 30 minutes organoid media was added. Cells were grown for 5 days with daily media changes and then digested using Liberase as before and passaged for expansion or assays. Organoid media recipe consists of Advanced DMEM/F12, 1× N-2, 1× B27,10 mM HEPES buffer,1× L-glutamine (2 mM), 1× penicillin–streptomycin (5,000 U/mL)(*47*).

### Cell culture and cell harvesting

Cell lines were grown in RPMI complete, 10% FBS with 1X Pen-strep. Cells were prepared for single-cell RNA sequencing by trypsin digestion for 10 minutes at 37C when harvested from in-vitro culture. For harvest of organoids and in-vivo xenograft tumors, cells were harvested by resection, chopping with a razor blade into a slurry, and then resuspension in 1 mg/mL Liberase TM. Suspensions were incubated shaking at 37C at 150 RPM for 45 minutes after which cells were resuspended in ACK buffer and incubated at room temperature for 10 minutes. Resuspension in 1X PBS was performed two times following this for wash and removal of digestion buffers. Cells were passed through a 70 micron mesh filter to arrive at single-cell suspensions.

### Cell suspensions and cell hashing

Cell suspensions were then hashed using HTO oligos or Totalseq-A antibodies. Cell suspensions were assigned unique oligo-HTO or antibody-HTOs at the point of experimentation. Cells were incubated with corresponding HTO conjugate for 20 minutes on ice. Cells were then washed 3X in 1X cold PBS and then recounted. Cells were then pooled together from multiple hashed suspensions such that there were equivalent numbers of cells from each suspension.

### Growth rate measurements, and cell line pooling

Cells were seeded in 6 well plates at 50k/well for a T0 time point. After 72 hours, cells were harvested by trypsin and counted to arrive at total cell numbers. Growth rates were calculated by the following formula: r = ln(Y t/Y 0)/t. Cell line pooling for GENEVA was performed by harvesting all cell lines simultaneously by trypsin digestion, washing in 1X PBS, counting, and inversely balancing against growth rate. Cells were seeded at specific numbers inverse to the growth rate into a single sterile tube and then mixed well before plating into separate wells in vitro, or injections into different mice for flank xenograft studies. All cell line pooling was performed in less than 2 hours total time to ensure healthy cell pools.

### Compound sources

Compounds were obtained from MedChemExpress, CaymanChem, and SelleckChem. Large batches of ARS-1620 were obtained with thanks from Dr. Kevan Shokat’s laboratory.

### Compound treatments

In vitro compound treatments were prepared by diluting the compound to final concentration in 1X RPMI complete, 10%FBS, pen-strep with no more than 1% DMSO in the final solution. Combination treatments were prepared in the same way. In vitro treatments received media changes with fresh compound dilutions every 48 hours. In vivo delivery was formulated as specified for the specific study and dosed by standard IP or PO dosing.

### GENEVA library preparation and sequencing

GENEVA libraries were prepared according to single-cell RNA sequencing Chromium kit instructions. Modifications were performed after the cDNA step in which the small fractions were harvested and then amplified in separate PCR reactions for enrichment of the hashtag reads, also as specified in Chromium kit. Sequencing was performed for the transcriptome and hashtag libraries at a 95%/5% ratio (transcriptome/hashtag). Sequencing targets were 25,000 reads/cell based on expected cell counts determined by number of cells loaded at input into the chromium kit.

### GENEVA data processing

#### Data pre-processing

Raw single-cell FASTQ files were used as input into the 10X *Cell Ranger* pipeline with default parameters. Briefly, reads were aligned to the hg38 transcriptome using *STAR* generating a filtered feature-barcode matrix and bam alignment file.

#### SNP detection and cell line identification

QuantSeq 3’RNA-Seq FASTQs for each cell line were first processed using *cutadapt* to remove adapter sequences. Processed 3’Seq reads were then aligned to the hg38 transcriptome using STAR. Bam files were then sorted and deduplicated using *UMI-tools dedup*, and used as inputs for variant calling using *bcftools mpileup* and *bcftools call* with default parameters. VCF files across all cell lines were merged using *bcftools merge*.

Pooled scRNA-Seq experiments were demultiplexed into individual cell lines using *Demuxlet* with the *CB* tag, setting the fixed mixing proportion, alpha, to 0.5. Demultiplexing of single cell barcodes into drug treatments using hashtag oligos (HTOs) was performed using *pymulti* from the scEasyMode python library (https://github.com/johnnyUCSF/scEasyMode)

#### Post-processing, doublet removal, quality checks and scoring

Demultiplexed scRNA-Seq count matrices were processed using *scanpy*. Counts were normalized (*normalize_per_cell*) and, log-transformed (*log1p*). Additionally, droplets predicted as doublets by *demuxlet* were removed. Normalized count matrices were used to calculate PCA, leiden clusters, and UMAP representations.

Cells with less than 1000 uniquely expressed genes (*n_genes < 1000*), greater than 15% mitochondrial fraction (*mito_frac > 0.15*) and less than 2000 counts (*n_counts < 2000)* were identified as apoptotic cells based on thresholds determined by https://pubmed.ncbi.nlm.nih.gov/31603610/

#### Differential expression analysis

The scanpy differential expression framework (Wilcoxon rank-sum test) was used to obtain log2 fold change (Log2FC) values between drug and control conditions. Log2FC values were then z-score normalized across cell lines and averaged to retrieve mean difference z-scores. Two-tailed t-tests for each gene were performed against a bootstrapped distribution of mean difference z-scores.

G12C specific expression was done by first calculating differential expression between DMSO and ARS-1620 treatment for each cell line and then taking the mean Log2FC difference between cell lines with G12C and non-G12C mutations. P-values were calculated using two-tailed t-tests between G12C and non-G12C groups.

#### Enrichment Analysis

iPAGE was used to conduct enrichment analyses using an information-theoretic framework as described in (*48*). Briefly, continuous expression vectors associated with every cell or gene, i.e. mitochondrial gene expression (Fig 3A) or differential expression Log2FC values (Fig S4D), are quantized into bins. Next, binary profile vectors are created, indicating the presence or absence of drug treatments or genes belonging to a geneset, in each of the bins. Then, iPAGE tests for the mutual information (MI) between these vectors and quantifies over and under-representation by modeling a hyper-geometric distribution.

#### Drug combination analysis

Drug combination efficacy, as measured by Bliss synergy scores, were evaluated by the Bliss independence model as described previously in Liu et al (2018). To obtain synergy covariates, first, DESeq2 (*49*) was used to obtain gene level linear model coefficients (β) using a cell line and drug treatment (single and dual) factor design, using un-normalized scRNA-Seq counts as inputs. For a given cell line, a gene’s synergy covariate was calculated as the difference between the sum of the coefficients derived from the single-drug models, i.e additive effect, and the coefficient of the dual-drug model. Formally, (β_*A*_ + β*_B_*) − β_*A*+*B*_, for drug A and drug B, in a given cell line. The average gene synergy covariate of a gene across all cell lines was taken and ranked based on t-statistic values generated from two-tailed t-test comparisons to a bootstrapped distribution.

#### Gini Index Transcriptome Quantification

To quantify this, we compared the relative abundance of cells from ARS-1620 treated and DMSO control samples within each of the 11 Leiden clusters for this cell line. We observed that the ARS-1620 treated cells largely reside in the neighboring clusters ‘11’ and ‘14’. We used the Gini coefficient to compare the unequal distribution of relative cell counts across the gene expression clusters. We then compared the observed Gini index to the null distribution of Gini coefficients calculated for randomly sampled clusters of equal size.

### Phenotypic assays

#### Survival assays

Cells were seeded in black clear bottom 96 well plates at 5K / well. Cells were allowed to adhere overnight and the next morning were treated with compounds at the specified concentrations in 100 uL of drug media after removal of seeding media. After 72 hours, cells were lysed by Cell-titer glo protocol for luminescence reading or incubated for 30 minutes in Calcein-AM after which fluorescent reading at FITC channels were performed for data collection.

#### Respirations assays

After acute treatment of cells for 2h with AMG510, cells were immediately taken into standard seahorse respirometer assay using the Seahorse XF Cell Mito Stress Test protocol: https://www.agilent.com/cs/library/usermanuals/public/XF_Cell_Mito_Stress_Test_Kit_User_Guide.pdf

#### Combination therapies *in vitro* and *in vivo*

In vitro compound treatments were prepared by diluting the compound to final concentration in 1X RPMI complete, 10%FBS, pen-strep with no more than 1% DMSO in the final solution. In vitro treatments received media changes with fresh compound dilutions every 48 hours. In vivo delivery was formulated as specified for the specific study and dosed by standard IP or PO dosing. Note that compounds were prepared as a single suspension for combinations in vivo and dosed as one daily treatment.

#### Mitochondrial membrane potential assay

Mitochondrial membrane potential was measured by incubating TMRE (Tetramethylrhodamine, Ethyl Ester, Perchlorate) at 1 uM for 30 minutes at 37C in base RPMI media. Cells were then washed twice in 1X PBS and trypsinized for readout by flow cytometry. Detection was performed at 488nm/575nm excitation/emission.

#### Mitochondrial ROS formation assay

Mitochondrial ROS was measured by incubating MitoSox-Red (from Thermo Fisher, M36008) at 0.5 uM for 30 minutes at 37C in RPMI base media. Cells were then washed twice in 1X PBS and trypsinized for readout by flow cytometry. Detection was performed at 488nm/575nm excitation/emission.

#### Caspase activation assay

Caspase activation was measured by incubating Caspase-3/7 detection reagent (Thermo Fisher, C10423) at 1 uM for 30 minutes at 37C in RPMI base media. Cells were then washed twice in 1X PBS and trypsinized for readout by flow cytometry. Detection was performed at 488nm/575nm excitation/emission.

#### Lipid peroxidation assay

Lipid Peroxidation was measured by incubating BODIPY 581/591 C11 detection reagent (Thermo Fisher, C10423) at 10 uM for 30 minutes at 37C in RPMI base media. Cells were then washed twice in 1X PBS and trypsinized for readout by flow cytometry. Detection was performed at 488nm/575nm excitation/emission.

#### In vivo experiments

### Animal studies

Animal studies were performed using NOD.Cg-Prkdcscid Il2rgtm1Wjl/SzJ (NSG) mice obtained from The Jackson laboratories. All mice were 8-12 week old females at the point of implantation. Animals were housed at the UCSF Helen Diller Preclinical Tumor Core for all experimentation in accordance with IACUC protocols.

#### Xenografts and tumor measurement

Tumors were injected subcutaneously at 100 uL each flank in a 1:1 ratio of 1X PBS to Matrigel. Cell suspensions were prepared at 20 million cells per milliliter and kept on ice during and before injection. Tumor measurements were performed by 3D caliper measurements.

#### In vivo drug treatment

In vivo drug treatment was performed by diluting compound in 100% DMSO, and then dilution of the compound subsequently by addition of Labrasol and distilled water for a final formulation of 10% DMSO, 70% labrasol, and 20% water. Dosing for all studies was performed by oral gavage at 100 uL with 5 on / 2 off daily dosing. Antimycin was separately dosed in 20% DMSO, 80% distilled water at 50 uL by intraperitoneal injection with 5 on / 2 off daily dosing.

## Data Availability

Meta-analysis of synthetic lethal CRISPR screen data for different compounds come from the following publicly available data: https://pubmed.ncbi.nlm.nih.gov/28985505/, https://www.science.org/doi/full/10.1126/science.abl5829, https://www.ncbi.nlm.nih.gov/pmc/articles/PMC6104643/?report=reader#!po=1.72414, https://journals.plos.org/plosbiology/article?id=10.1371/journal.pbio.2004624#sec039, https://journals.plos.org/plosgenetics/article?id=10.1371/journal.pgen.1008057#sec034, https://pubmed.ncbi.nlm.nih.gov/31138768/

The data generated in this study is available in the Gene Expression Omnibus (GEO) under the accession number GSE283335.

## Code Availability

The code used in this study is available in the Gene Expression Omnibus (GEO) under the accession number GSE283335.

## Supplemental Figures

**Supplemental Fig 1.**
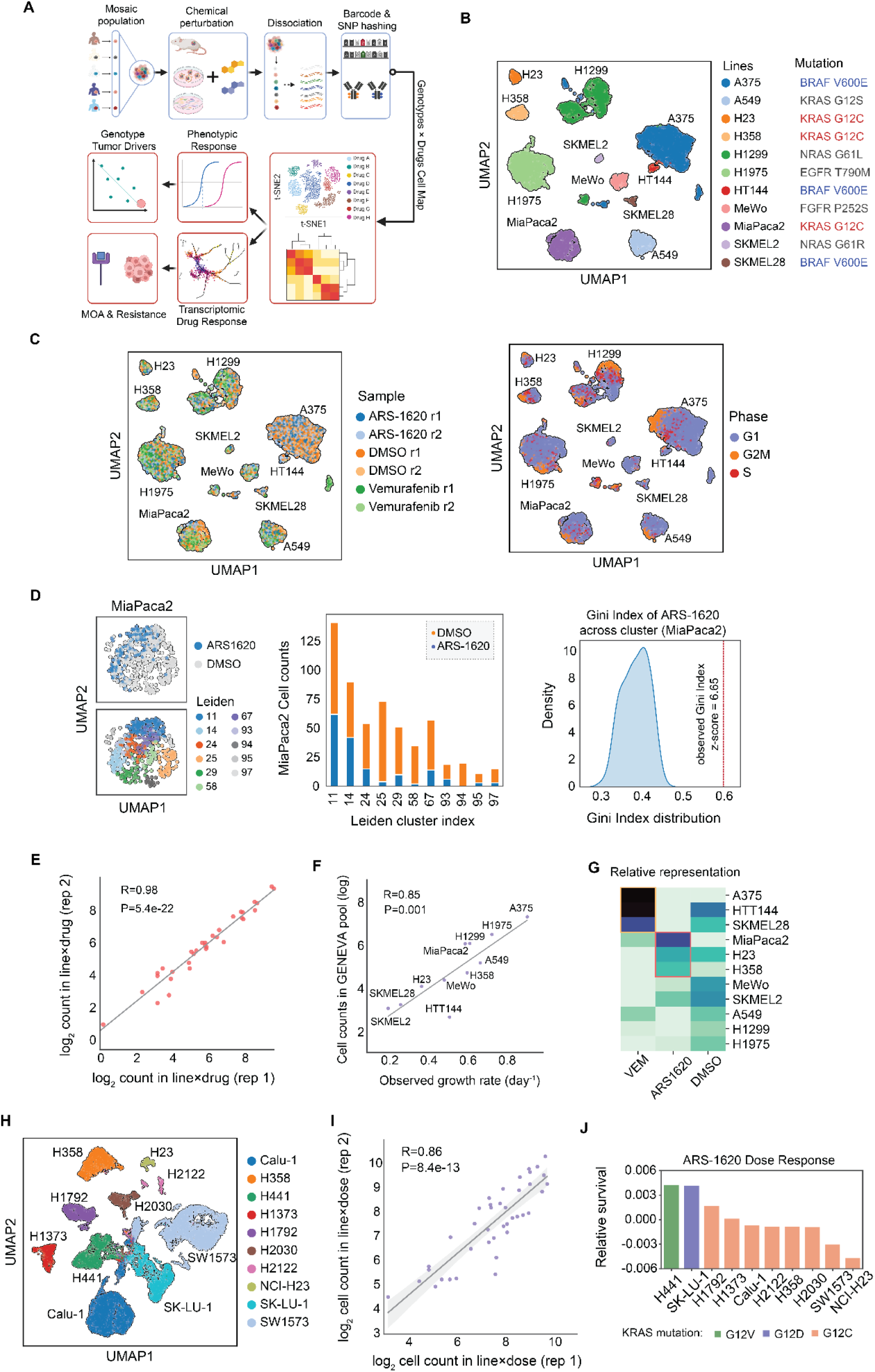
GENEVA schematic. **(A)** Conceptual overview of GENEVA as a multiplexed method for rich phenotyping of diverse populations. Mechanisms of drug efficacy and escape are elucidated using a mosaic tumor model approach. **(B)** UMAP projection of a GENEVA pool composed of heterogeneous cell lines from different genetic driver backgrounds. Clusters are colored by cell line identity, **(C)** drug treatment condition, and cell cycle phase. **(D)** UMAP projection of KRAS G12C mutant cell line MIA PaCa-2 colored by drug treatment condition and leiden sub-clustering (left). Cells binned by cluster representation across drug treatment and vehicle groups (center). Gini index distribution compared for drug and vehicle conditions across clusters (right). **(E)** Log_2_ of cell counts across two independent replicates showing concordance of cell counts after mosaic pool growth. **(F)** Comparison of Log_2_ cell counts observed with growth rates measured in vitro. **(G)** Heatmap of relative drug sensitivity in treatment conditions normalized on a per cell line basis from GENEVA cell counts. Dark blue and white indicate lower and higher relative representation respectively. **(H)** UMAP projection of a GENEVA pool with a cell line composition focused on KRAS.G12 mutant alleles. Clusters are colored by cell line identity. **(I)** Log_2_ of cell counts from focused KRAS G12* pool across two independent replicates. **(J)** Relative survival of cell pools normalized to vehicle pools for each cell line. Cell lines are colored by the identity of KRAS G12* mutation.

**Supplemental Fig 2.**
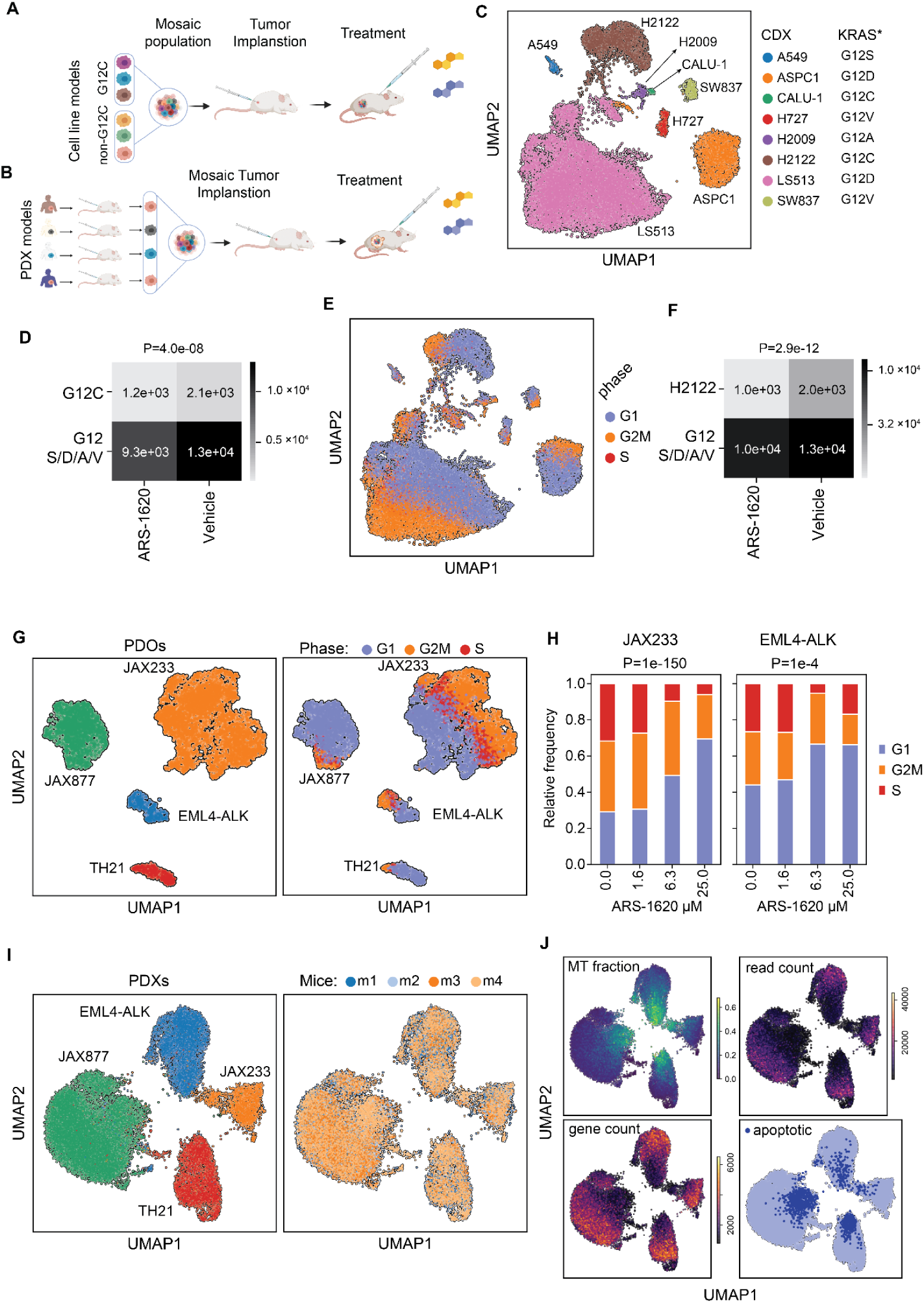
*In vivo* applications of GENEVA. Schematic of GENEVA for cell line derived xenografts. Mosaic populations are formed by pooling cell lines of interest followed by implantation in mice. Here, we have performed subcutaneous injections, however, for cell lines of similar origin orthotopic injections can also be performed. Once the mosaic tumors have grown, the xenografted mice can be treated with the drugs of interest for several days prior to tumor extraction. Once extracted, the tumors are dissociated and subjected to mouse cell removal and single cell profiling. Two focused experiments are illustrated using **(A)** cell line-derived xenograft models and **(B)** patient-derived xenograft models. **(C)** Cell line-derived xenograft GENEVA pool visualized as UMAP colored by cell line identity. (**D)** Contingency table showing significance of cell line counts in KRAS G12C and non-G12C across ARS-1620 and vehicle conditions. Χ² was used to calculate P-value. **(E)** CDX pool UMAP colored by cell cycle phase. **(F)** Contingency table showing significance of cell line counts in H2122 vs non KRAS G12C lines across ARS-1620 and vehicle conditions. Χ² was used to calculate P. **(G)** UMAP of PDOs grown in GENEVA mosaic colored by PDX identity (left) and cell cycle phase (right) after treatment with multiple doses of ARS-1620. **(H)** G1 arrest of PDXs across increasing ARS-1620 dose shown as proportion of cells in each cell cycle phase. Χ² was used to calculate P. **(I)** PDX *in vivo* GENEVA pool visualized as UMAP. Cells are colored by PDX identity and mouse replicates. **(J)** Cells are further colored by several metrics calculated on an individual cell basis: reads from mitochondrial fraction, total read count, total genes detected, and apoptotic cell state. See detailed methods for scoring methods for single-cell transcriptomes.

**Supplemental Fig 3.**
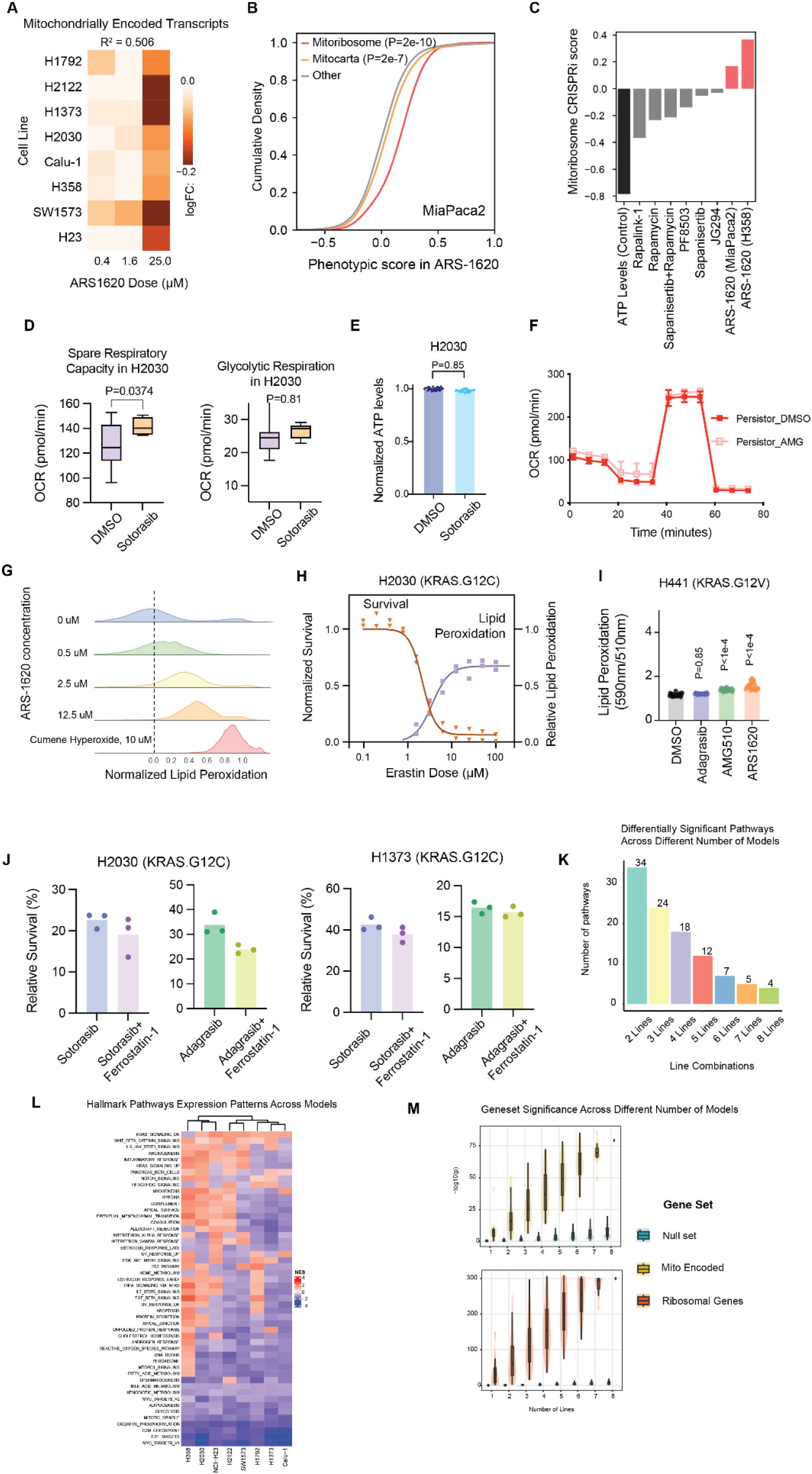
Effects of G12Ci on mitochondrial and lipid peroxidation pathways. **(A)** Mitochondrial transcripts across GENEVA pools at escalating doses of ARS-1620, normalized to vehicle-treated pools. Colors shown reflect Log_2_ fold-change of mitochondrial transcripts for cell lines at different doses relative to vehicle. R^2^ was calculated by treating log_2_FC values as the response variable, cell lines as replicates, and dose as the independent variable. **(B)** Genome-wide screen data from KRAS G12C mutant MIA PaCa-2 lines in a synthetic lethal screen with ARS-1620 grouped by MitoCarta, Mitoribosome, and all other genes. Increase in phenotypic score indicates a buffering or rescuing effect on cell survival in a drug treatment context. Genes were treated as replicates within a geneset (Mitoribosome, Mitocarta) and compared using a two sample *t*-test against the majority of other genes in the genome-wide screen data (ie non-Mitoribosomal, non-Mitocarta). **(C)** Median phenotypic score for mitoribosomal genes in multiple genetic screens. All shown values are significantly different from the background values based on Mann-Whitney *U*-test. **(D)** Spare respiratory and glycolytic respiration in H2030 after acute treatment with Sotorasib as measured by OCR. P-values are obtained using a two sample *t*-test. **(E)** ATP levels as measured by Cell Titer Glo after acute treatment with Sotorasib at 2h, 500 nM. P-values were obtained using a two sample *t*-test. **(F)** Persistor population after long-term ARS-1620 treatment is treated with acute Sotorasib and assayed by Seahorse respirometer. No significant difference observed by a two sample t-test across all respiratory compartments. **(G)** Lipid Peroxidation measurement of cells by flow cytometry after dose titration of ARS-1620. Lipid peroxidation measurements were performed at 24 h post-treatment using BODIPY C11 reagent and compared to Cumene Hyperoxide, a known rapid inducer of lipid peroxidation as a control. **(H)** Relative cell survival and lipid peroxidation levels dose-response to Erastin. Survival was measured using CTG at 72 hours and lipid peroxidation with BODIPY C11 at 24 hours. Left axis shows normalized survival and the right axis shows normalized lipid peroxidation. **(I)** Lipid peroxidation values after treatment with three KRAS G12C inhibitors compared to a vehicle in H441 (KRAS G12V). P-values are obtained using a two sample t-test against vehicle (DMSO). **(J)** Cell survival proportions with Sotorasib and Adagrasib combination dosing with Ferrostatin-1 in two KRAS G12C cell lines (H2030 and H1373) and one non-G12C cell line (H441). No rescue was observed with addition of Ferrostatin-1 pretreatment for 24 hours. **(K)** Downsampling of multiple cell lines from MOSAIC data and significant geneset number as a function of the number of models included. Hallmark pathways were identified in individual cell line comparisons using a significance threshold of padj < 0.05 computed using GSEA. Intersections of these pathways were taken for all combinations of cell lines and the set size of the union of these intersections was plotted. **(L)** Normalized Enrichment Scores (NES) for GENEVA cell lines after KRAS G12C inhibitor treatment. NES scores were calculated by gene set enrichment analysis (GSEA) on z-score normalized differential expression values for each cell line, individually. The heatmap shows NES for significantly enriched Hallmark pathways (padj < 0.05) in at least one cell line **(M)** Downsampling of multiple cell lines from MOSAIC and individual geneset examination for Mitochondria resident genes and Ribosomal genes across p-values. P-values for mitochondrial resident and ribosomal gene sets were calculated in all combinations of cell lines from MOSAIC by combining individual p-values using Fisher’s method. This was compared to p-values calculated in the same way from 100 iterations of randomly sampled null gene sets of 100 genes.

**Supplemental Fig 4.**
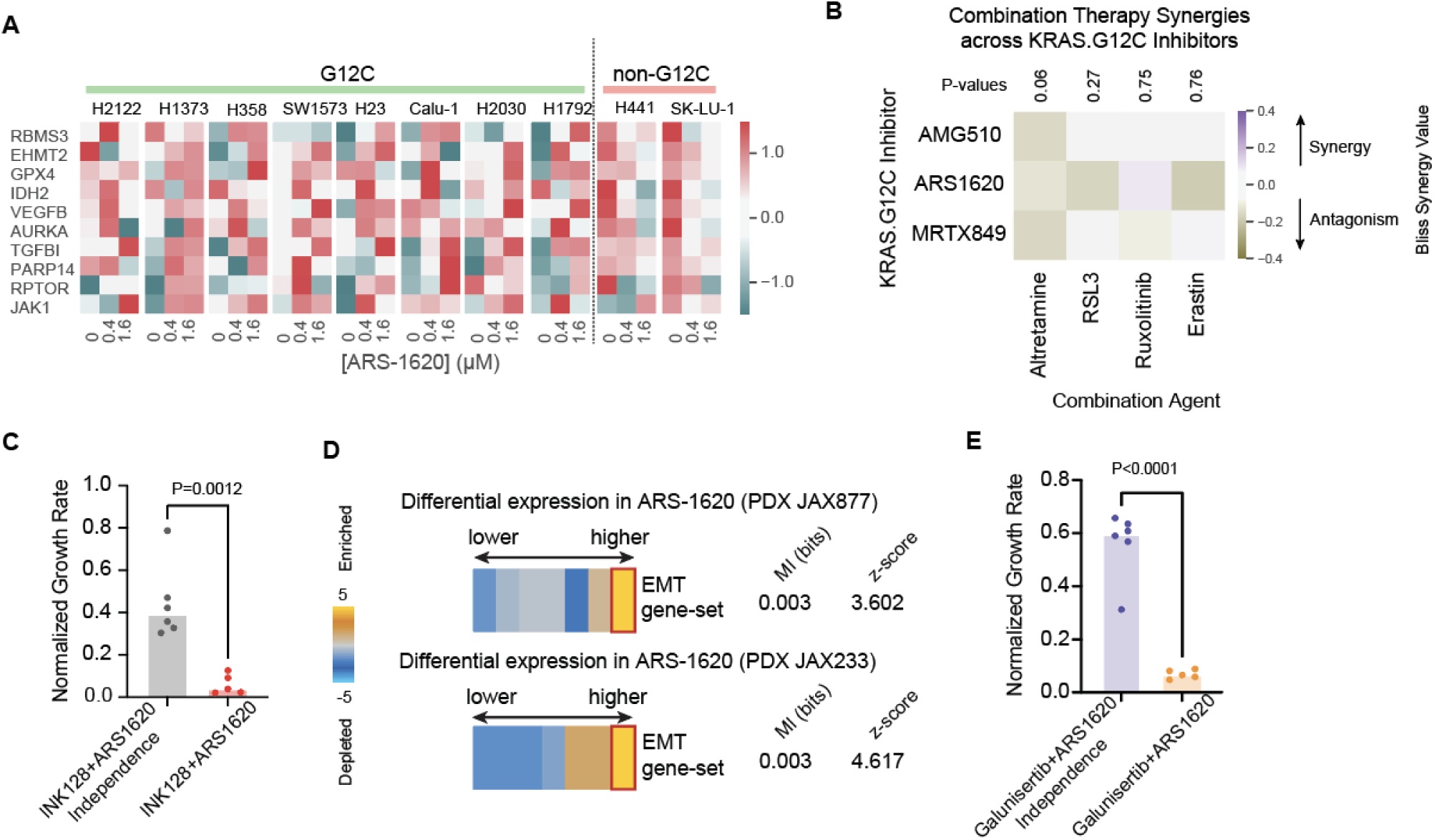
Discovery of mechanisms of resistance to KRAS G12C inhibition. **(A)** Heatmap of gene expression of various G12C specific mechanisms across individual cell lines and ARS-1620 dose. **(B)** Bliss synergy of combination of KRAS G12C inhibitors and three ferroptosis inducers and Ruxolitinib. *P-values* were calculated by two-tailed t-test of replicates specific to a dual-agent combination compared against a null distribution estimated single agent activities. **(C)** *In vivo* combination studies of INK128+ARS-1620 visualized as the expected additive effect on growth rate of tumors and the observed *in vivo* growth rate. P-value obtained using a two sample *t*-test. **(D)** EMT upregulation in PDX GENEVA pools for two KRAS G12C mutant PDX after longitudinal drug treatment. The EMT gene-set is significantly enriched in the top-most gene expression bin for both JAX877 and JAX233 PDX models. Shown are the enrichment (gold) and depletion (blue) patterns of this gene-set across the differential expression bins, as well as their association mutual information (MI) and *z*-score. **(E)** *In vivo* combination studies of INK128+ARS-1620 visualized the expected additive effect on growth rate of tumors and the observed *in vivo* growth rate. P-value are obtained using a two sample *t*-test.

**Supplemental Fig 5.**
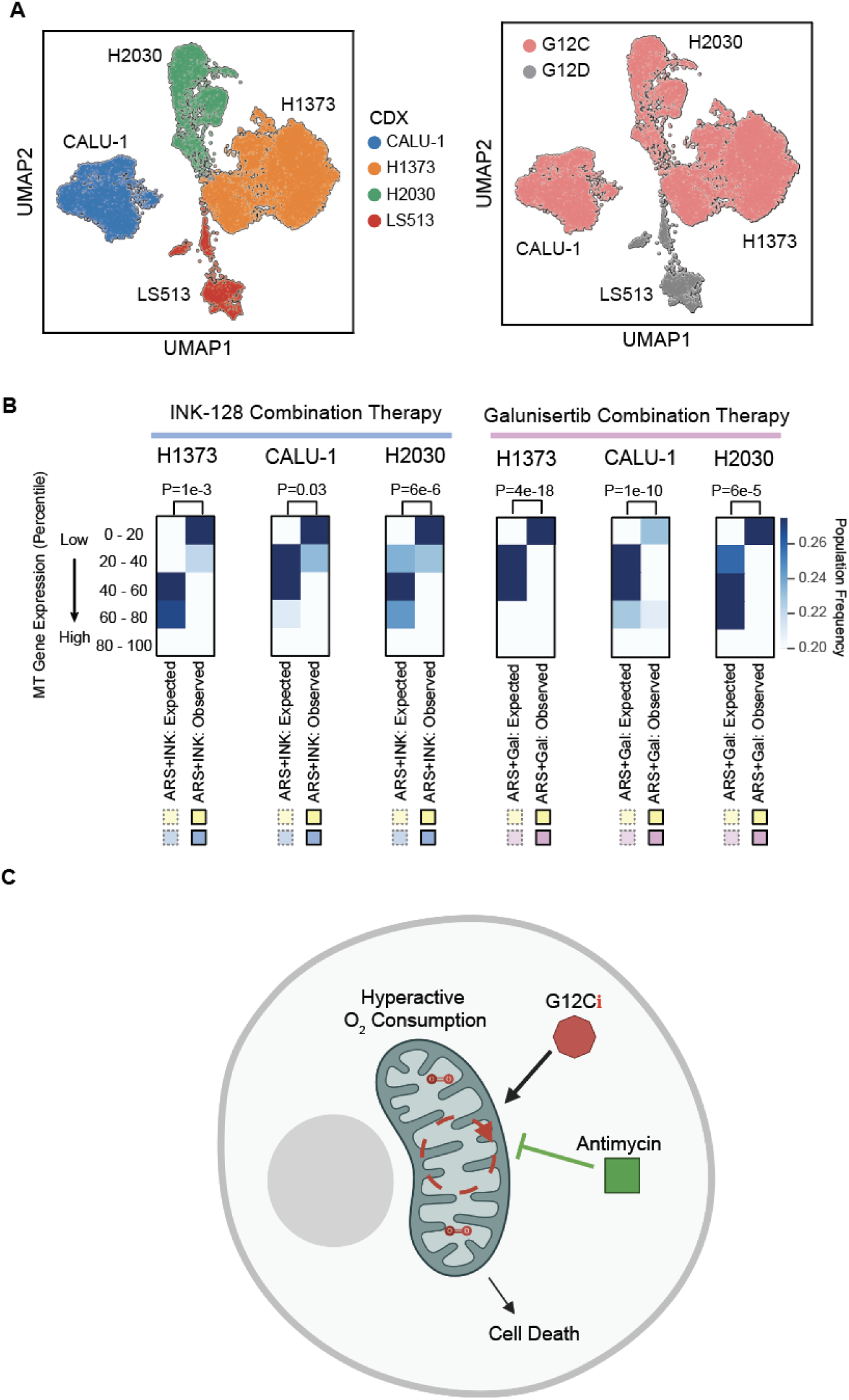
GENEVA enables gene level profiling of *in vivo* drug combinations. **(A)** UMAP visualization of an *in vivo* GENEVA pool for combination therapy studies colored by cell line identity (left) and G12 mutant allele (right). **(B)** INK128 and Galunisertib with ARS-1620 gene level synergy model. Blue squares indicate enrichment of cells across bins of mitochondrial expression. Gene level synergistic decrease in mitochondrial genes was measured by two tailed t-test. **(C)** Model of KRAS G12C pharmacological inhibition’s effect on mitochondrial oxygen turnover leading to cell death.

